# Fluorescence Multiplexing with Spectral Imaging and Combinatorics

**DOI:** 10.1101/361790

**Authors:** Hadassa Y. Holzapfel, Alan D. Stern, Mehdi Bouhaddou, Catilin M. Anglin, Danielle Putur, Sarah Comer, Marc R. Birtwistle

## Abstract

Ultraviolet-to-infrared fluorescence is a versatile and accessible assay modality, but is notoriously hard to multiplex due to overlap of wide emission spectra. We present an approach for fluorescence **<u>m</u>**ultiplexing **<u>u</u>**sing **<u>s</u>**pectral **<u>i</u>**maging and **<u>c</u>**ombinatorics (MuSIC). MuSIC consists of creating new independent probes from covalently-linked combinations of individual fluorophores, leveraging the wide palette of currently available probes with the mathematical power of combinatorics. Probe levels in a mixture can be inferred from spectral emission scanning data. Theory and simulations suggest MuSIC can increase fluorescence multiplexing ~4-5 fold using currently available dyes and measurement tools. Experimental proof-of-principle demonstrates robust demultiplexing of nine solution-based probes using ~25% of the available excitation wavelength window (380-480 nm), consistent with theory. The increasing prevalence of white lasers, angle filter-based wavelength scanning, and large, sensitive multi-anode photo-multiplier tubes make acquisition of such MuSIC-compatible datasets increasingly attainable.

## INTRODUCTION

Fluorescence in the UV to infrared range is one of the most widely-used and easily accessible quantitative and qualitative assay modalities across the life and physical sciences^1^. Yet, fluorescence is notoriously hard to multiplex; that is, to measure multiple analytes simultaneously in a mixture. Typical fluorescence multiplexing is routinely limited to about four colors, each corresponding to a single measurement^2^. For example, one of the arguably most multiplexed and data dense experimental modalities—Illumina “next-generation” deep DNA sequencing, relies on such four-color imaging; one for each DNA base^3^. This four-color standard is the case when fluorescence emission is collected via broad-banded filters, as opposed to the entire emission spectra.

When so-called hyper-spectral, or fluorescence emission scanning is employed along with linear unmixing^4,5^, measurement of up to seven analytes is possible^6^. Cycles of staining tumor sections with fluorophore-labeled antibodies, coupled with chemical inactivation and multiple rounds of staining has reported to analyze 61 antigens^7^. A similar principle has been applied without the use of a proprietary instrument to produce cyclic immunofluorescence that uses repeated rounds of four color imaging for ~25 analytes^8^.

Specific assay instantiations that separate analytes in a variety of ways have also been able to reach higher multiplexing capabilities. For example, super-resolution imaging combined with *in situ* hybridization and combinatorial labeling used fluorescence to measure 32 nucleic acids in single yeast cells^9^. The Luminex xMAP system can multiplex ~40 analytes separated by specific beads^10^. Segregating fluorophores by individual bacterium can multiplex ~ 28 different strains using “CLASI-FISH”^11^. Alternatives to fluorescence are also of course many; for example mass cytometry which measures levels of ~30 specific isotope tags as opposed to fluorophores^12,13^.

Despite these advances, there remains yet to be reported, to our knowledge, a fluorescence-based technology that simultaneously can demultiplex more than 4-7 analytes within a mixture. Such an ability may have widespread impact, due to the prevalence, sensitivity and versatility of fluorescence as a measurement tool. Here, we report such an advance which we term **<u>m</u>**ultiplexing **<u>u</u>**sing **<u>s</u>**pectral **<u>i</u>**maging and **<u>c</u>**ombinatorics (MuSIC). MuSIC works by creating covalent combinations of existing fluorophores and measuring fluorescence emission spectra of their mixtures. We first describe the theoretical basis for MuSIC, and then through simulation studies explore potential limits of the approach. Finally, we experimentally demonstrate the feasibility of MuSIC to measure the levels of nine different fluorescent probes in a mixture using only ~25% of the available spectral window of fluorescence excitation (380-480 nm), supporting a potential 5-fold increase in fluorescence multiplexing ability. The advent and accessibility of white lasers, angle filter-based emission wavelength scanning, and large, sensitive multi-anode photo-multiplier tubes make acquisition of such MuSIC-compatible datasets increasingly attainable.

## RESULTS AND DISCUSSION

### Theory

Fluorescence emission follows principles of linear superposition^2^. Therefore, the emission spectra of a mixture of fluorophores can be cast as the sum of its component parts with a matrix equation (Fig. 1A)

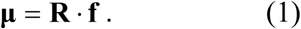

**Figure 1.**
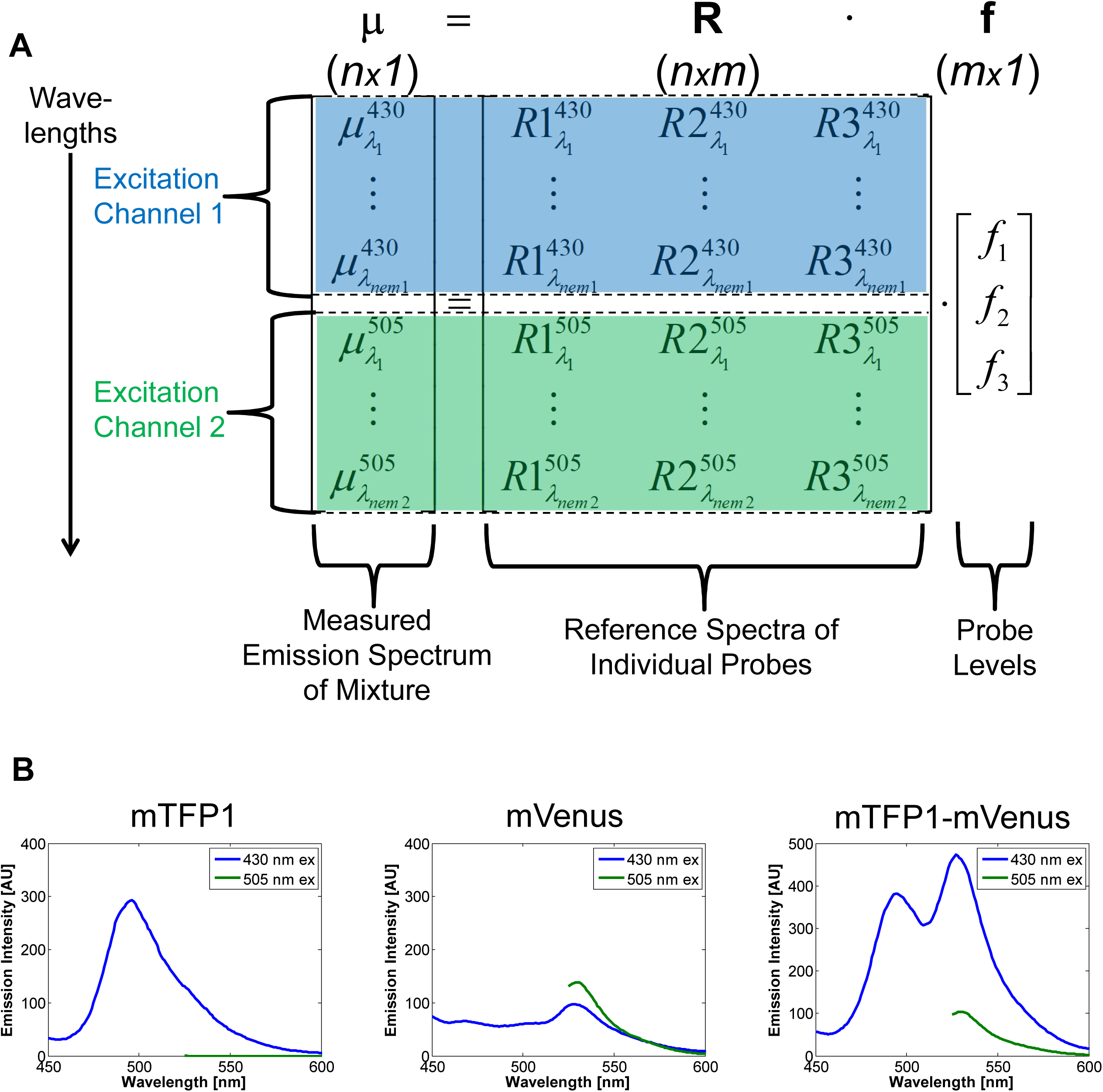
Theoretical Basis for Multiplexing using Spectral Imaging and Combinatorics (MuSIC). **A.** Example arrangement of data for a three probe (*m*=*3*) setup in terms of the linear unmixing equation. Emission spectra data (μ) of a mixture are arranged vertically, stacked by emission wavelength and excitation channel (indicated by color and background highlighting). Each column of the reference matrix is the emission spectra of a probe in isolation, arranged in the same way. **B.** Example data for a three probe setup involving a teal fluorescent protein (mTFP1), a yellow fluorescent protein (mVenus), and their covalent fusion (mTFP1-mVenus). Two excitation channels are used, 430 and 505 nm, and fluorescence emission spectra are measured.

Here, **μ** is an *n*-by-*1* vector of measured fluorescence emission intensities at *n* emission wavelength/excitation channel combinations, **R** is the *n*-by-*m* matrix of reference emission intensity spectra for *m* individual fluorescent probes aligned in columns (which could include a column for background fluorescence), and **f** is an *m*-by-*1* vector containing the relative levels of the *m* individual probes. The reference spectra correspond to those of each individual probe in isolation. Note that this equation also can account for multiple *n_ex_* excitation channels (Fig. 1A).

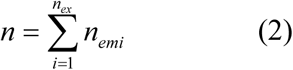

where *i* denotes excitation channel index, and *n_emi_* is the number of emission wavelengths measured in that excitation channel. In this case, the rows in every column of the reference matrix **R** must be arranged in the same order of excitation channels and wavelengths, along with the measurements **μ** (Fig. 1A).

Solving Eq. 1 for **f** to infer the relative levels of *m* individual probes is called “linear unmixing”^4,5^. Mathematically, solving for **f** requires the rank of the matrix **R** to be greater than or equal to *m*. By increasing the rank of **R**, one increases the number of individual component levels *m* that can be independently estimated from fluorescence emission spectra measurements. A typical way to increase the rank of **R** is to use multiple excitation channels, which is the intuitive basis for traditional multi-color imaging. Yet, increasing the number of excitation channels does not guarantee increasing the rank of **R**, because redundant information could be added. For example, exciting yellow fluorescent protein (YFP) variants with 505 and 510 nm light would usually not increase the rank of **R** because they are excited in a similar manner by both of these excitation wavelengths.

Multiplexing with Spectral Imaging and Combinatorics (MuSIC) works by using covalently-linked combinations of fluorophores to add columns to **R** which increase its rank. Each new fluorophore combination has a new column in **R**, and if it increases the rank of **R** by one, then in theory its levels can be estimated through linear unmixing (we use simulation below to explore this more practically with added noise). Consider here a simplistic illustration with experimental data from teal fluorescent protein (mTFP1) and YFP (mVenus) (Fig. 1B). If one excites at 430 nm and 505 nm, then mTFP1 and mVenus emission are largely separated by independent excitation, and one can quantify the levels of mTFP1 and mVenus in a mixture, a standard two-color experiment. However, in the spectral emissions from both channels, there is “room to carry” more information, and in particular the red-shifted portion of 430 nm excitation channel. Because the excitation spectrum of mVenus overlaps with the emission spectrum of mTFP1, they exhibit fluorescence resonance energy transfer (FRET) when in close proximity. By including an mTFP1-mVenus fusion in the experiment, the acceptor mVenus emission becomes strongly visible in the 430 nm channel by FRET (Fig. 1B, far right panel). This increases the rank of **R** by one, and allows quantification of mTFP1, mVenus, and mTFP1-mVenus levels in a mixture.

This analysis suggests to us that the creation of a new MuSIC fluorescence probe requires that (i) there is sufficient FRET to allow observable fluorescence emission of the acceptor in a new excitation channel and (ii) the resulting emission spectra of the new combination fusion probe is sufficiently distinct from all the other probes in at least one excitation channel. We use these guidelines in the subsequent simulation studies to explore the potential limits of this line of reasoning and more precisely define these sufficiency criteria.

### Simulation Studies to Explore Limits and Potential of MuSIC

The above theoretical considerations suggest that MuSIC may offer large increases in fluorescence multiplexing capabilities. How many probes might be multiplexed and their levels estimated simultaneously from a mixture? We performed simulation studies to give insight into these questions (Fig. 2A). Specifically, we considered 16 individual fluorescent proteins (FPs): EBFP2^14^, mTagBFP2^15^, mT-Sapphire^16^, mAmetrine^17^, mCerulean3^18^, mTFP1^19^, LSSmOrange^20^, EGFP^21^, TagYFP^22^, mPapaya1^23^, mOrange2^24^, mRuby2^25^, TagRFP-T^24^, mKate2^26^, mCardinal^27^, and iRFP^28^. We selected these to span the UV to IR spectrum, for reported photostability, and approximately similar brightness (although this last task is reasonably difficult). We hypothesized that having similar brightness levels would help to increase dynamic range. It may be possible to equalize brightness better across probes by considering integer multiples of FPs; for example a tandem dimer should roughly double the effective brightness.

**Figure 2.**
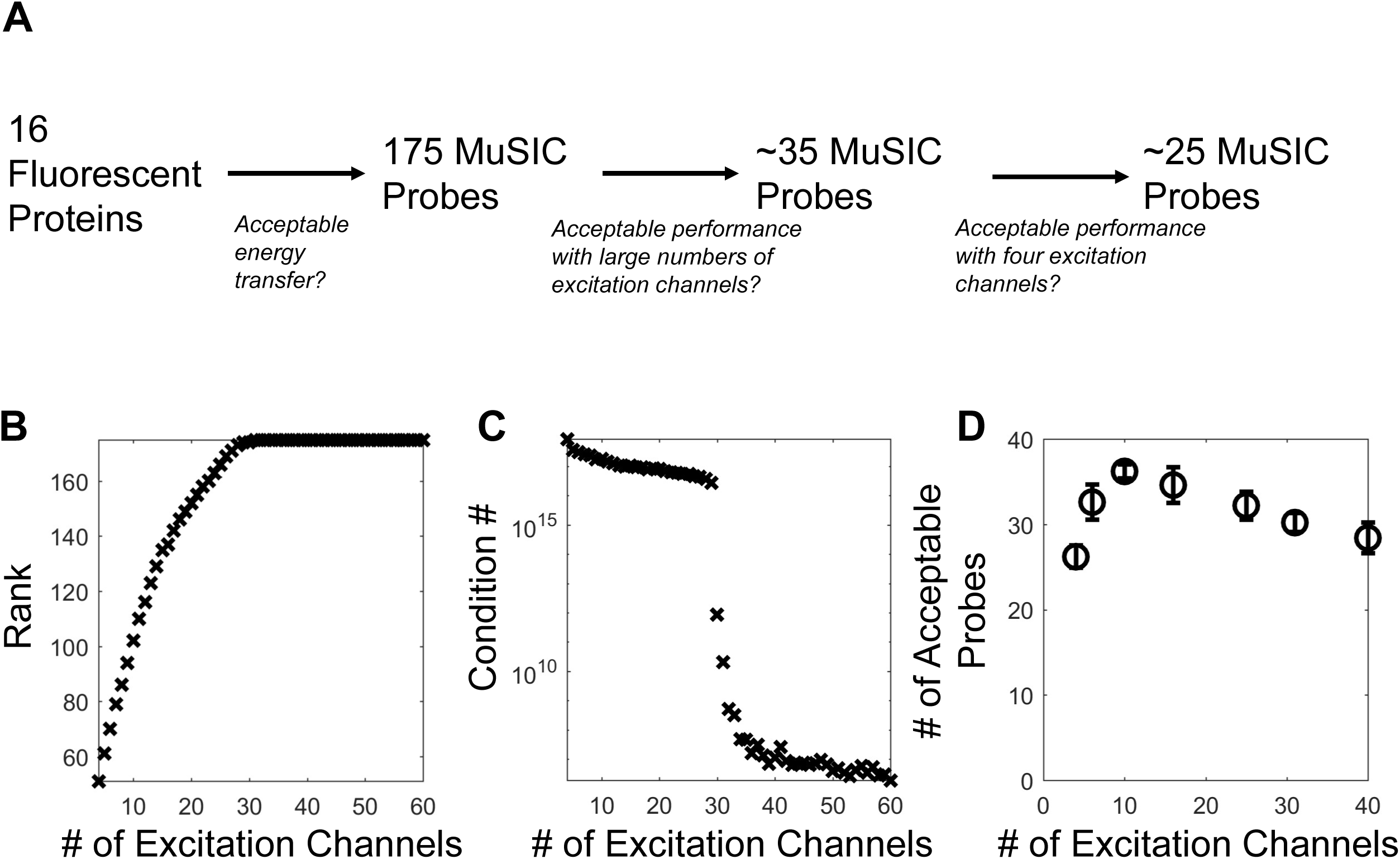
Simulation Studies for the Potential and Limits of MuSIC. **A.** Summary of how 16 considered fluorescent proteins are converted into 175 putative MuSIC probes, of which ~35 have acceptable quantitative behavior using ~10 excitation channels, and ~25 using four excitation channels. **B.** Rank of the reference matrix containing all 175 potential MuSIC probes as a function of the # of excitation channels. Full rank is achieved at 31 excitation channels. **C.** Condition number (log scale) of the reference matrix containing all 175 potential MuSIC probes as a function of the # of excitation channels. **D.** The number of probes that have a correlation coefficient greater than 0.7 after reducing the number of probes from the original 175, as a function of the # of excitation channels. Error bars denote standard deviation across five probe reduction simulations.

The first aspect of this simulation study was to consider how to combine the individual FPs. Bimolecular FRET is common^29^, and trimolecular FRET less so, but has been reported^30^. We therefore exhaustively considered single FPs, dimers and trimers, but filtered all dimers and trimers where FRET efficiency was expected to be < 0.2 (based on calculated spectral overlap integral—see Methods). This gave rise to 175 probes that could potentially be quantified from perfect noise free measurements so long as **R** is of full rank (see Supplementary Table 1). However, determining the rank of **R** requires selecting excitation wavelengths.

We first considered a scenario where using a large number of evenly spaced excitation channels between 350 and 700 nm was feasible. We varied the number of excitation channels from four to 60, estimated the relative excitation strength of each probe at each excitation wavelength (from known excitation spectra), and summed the calculated emission intensity in 1 nm increments from 300 to 850 nm (based on the excitation strength, predicted FRET efficiencies, reduced FRET efficiency from direct acceptor excitation, and FP brightnesses—see Methods). There are diminishing returns past 31 excitation channels, where **R** saturates at a full rank of 175 (Fig. 2B). The condition number is a metric that can be thought of quantifying practical rank of a matrix, where lower numbers indicate a better ability to solve the linear unmixing problem in Eq. 1. The condition number also starts to decrease sharply around the same number of excitation channels (Fig. 2C). However, its magnitude suggests that unmixing performance may be inadequate in terms of *%* error; values ~10^7^ indicate a likely ill-conditioned matrix, and this large value decreases marginally with increasing number of excitation channels.

We next sought to identify how many probe levels might be reliably quantified using a large number of excitation channels (40), and also how that number of probes changes as the number of excitation channels is reduced down to a more typical four. We simulated multi-spectral measurements 20 times by sampling probe levels between 0 and 1000 relative concentration units, calculating the expected emission spectra (as above), adding noise to those spectra (similar to what is measured in below experiments), and then unmixing to estimate the probe levels. We quantified performance with a Pearson correlation coefficient *ρ* between the known, randomly sampled levels and estimated, unmixed levels for each of the 175 (or fewer) probes. We progressively eliminated those probes with the lowest correlation coefficient until all probes could be reliably quantified over 3 orders of the sampled concentration magnitude with *ρ*>0.7 in simulations.

In the case of 40 excitation channels, first we found that the same sets of probes were not recovered in independent simulation runs. This is because adding noise to simulated fluorescence emission spectra data is random, which causes random probes to have the worst correlation coefficient during the removal process. Therefore, we simulated probe removal five independent times for each number of excitation channels considered. We could not pinpoint discernable patterns for which probes were included across multiple simulation runs (full results in Supplementary Table S2); single, double and triple probes were prevalent, across the spectrum of available colors. This led us to hypothesize that the number of probes was much more important than probe identity, and that performance would likely have to be assessed experimentally on a probe by probe basis.

For 40 excitation channels, we found roughly 30 probes could be reliably quantified (Fig. 2D). Surprisingly, as the number of excitation channels was reduced, this number stayed constant or even slightly increased, all the way to 10 excitation channels where the number of probes was ~35. We speculated that this increase may be due to high quality probes being less likely to be removed during the culling process, although the exact reasons were difficult to pinpoint from the simulation data. With the advent and affordability of white lasers^31^, angle-tuned filters for wavelength scanning^32,33^, and large, sensitive multi-anode photo-multiplier tubes^34^, and an ever-increasing number of highly photostable fluorophores, such large excitation channel experiments may become or are already feasible. Below 10 excitation channels, the number of reliable probes decreases, although not drastically. With the current standard of four excitation channels, simulations suggest that approximately 25 MuSIC probes can be reliably quantified in a mixture. These simulation results suggest that MuSIC may provide a ~6-fold increase over a standard four color experiment, and up to ~8-9 fold if 10 excitation channels are used.

### Experimental Proof-of-Principal

We wanted to test MuSIC experimentally. Rather than fully expand to the entire spectrum of UV to infrared, we focused on a reduced range of ~25% of the available excitation spectrum from 380 to 480 nm, using the simulation studies above as a guideline for emission spectra every 1 nm, and 10 excitation channels. We reasoned that results here could be expanded and scaled subsequently after determining what caveats and limitations are revealed by reduction to practice that were not uncovered through the theory and simulation studies. This focused us on nine individual or combination probes that we created with fluorescent proteins (FPs) (Fig. 3A). We cloned, expressed and purified these proteins, (*E. coli*, His tag), and measured the reference spectra of each to verify (i) identity and (ii) appreciable FRET efficiency (Fig. S1). Next, we created 48 different mixture samples from these nine individual probes spanning 2-way probe combinations to all probes present. We prepared these mixtures in triplicate. We measured the emission spectra of these mixtures in 1 nm increments from 10 equally spaced excitation channels from 380 nm to 480 nm. From these spectral emission scanning data, we solved Eq. 1 to estimate the probe levels in each mixture. These “inferred levels” from estimates are compared to the “actual levels” for analysis.

**Figure 3.**
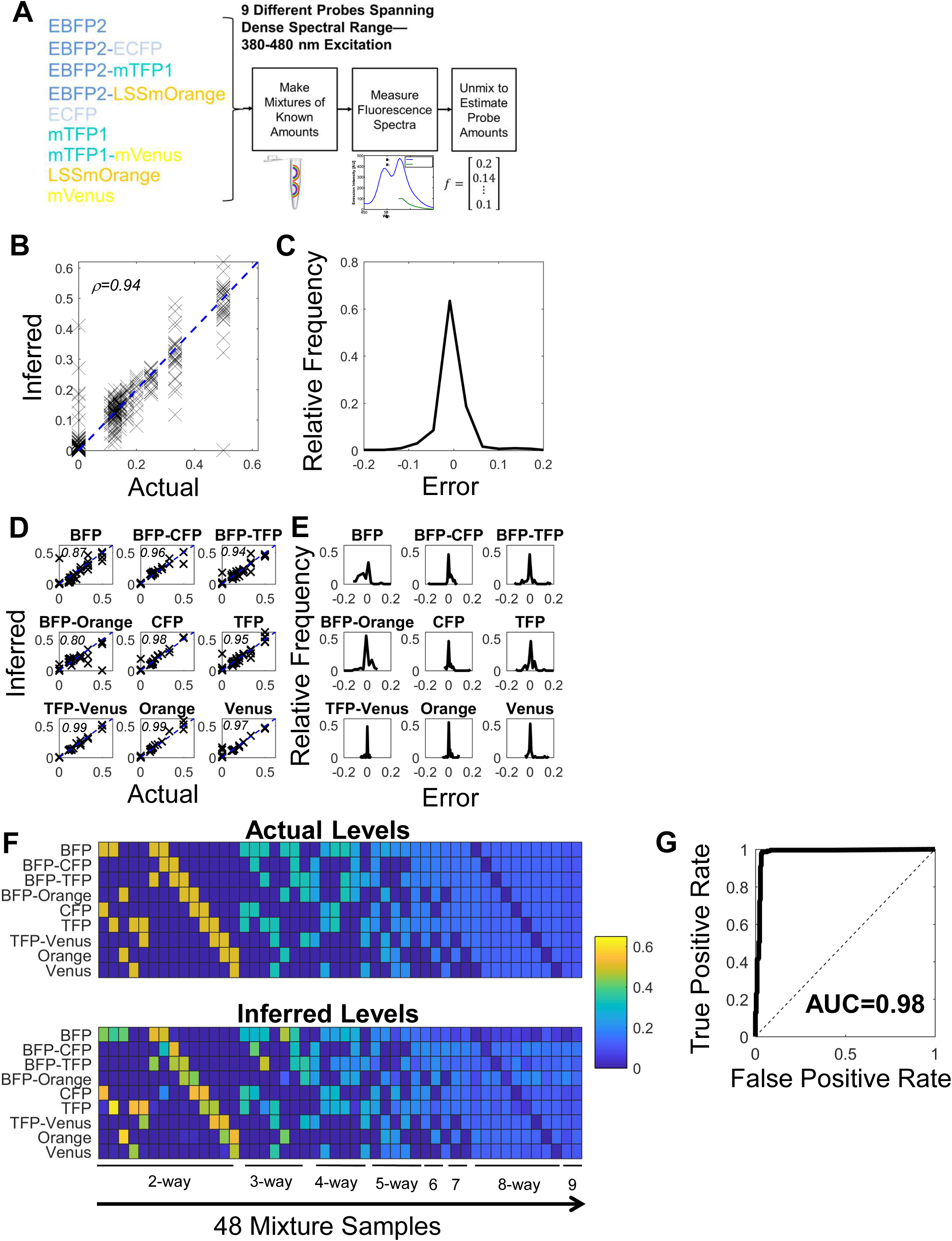
Experimental Evaluation of MuSIC. **A.** Experimental design. Nine different MuSIC probes were constructed from five fluorescent proteins as indicated. Excitation wavelengths were limited to between 380 and 480 nm. The pure probes were combined into mixtures with known amounts, their emission spectra were measured, and then their levels were estimated via unmixing. These inferred levels were compared to the actual, known levels in the mixture. **B.** Aggregate quantitative agreement between actual and inferred levels across mixtures and probes. Dashed blue line is x=y; Pearson’s correlation coefficient is shown. **C.** Histogram of errors, defined as the difference between actual and inferred probe levels, across mixtures and probes. **D-E.** Analogous plots as in B-C, respectively, segregated by probe. In D, text in upper left is Pearson’s correlation coefficient. **F.** Heatmaps of quantitative agreement between actual (top) and inferred (below) levels broken down by probe (vertical) and mixture (horizontal). Text below refers to the type of mixture; 2-way means two probes were included in the mixture, 3-way, three, and so on. Colorbar indicates relative probe levels. **G.** Receiver-operator characteristic (ROC) curve for binary classification of probe presence or absence across the 48 mixtures. The inferred probe level was compared to a threshold for classification as present or absent. This threshold was varied to generate the ROC curve, and was uniformly applied across samples and probes. AUC: area under the curve.

We first evaluated quantitative comparison between actual and inferred levels across all 48 samples and probes in aggregate (Fig. 3B-C). This analysis revealed reasonable agreement with most samples falling on or very close to the x=y line (black dashes in Fig. 3B; Pearson’s *ρ*=0.94), with only a few outliers away from this curve, and largely unbiased and symmetric error. We parsed these analyses by probe (Fig. 3D-E), which revealed that not all probes performed equally. For example, BFP and BFP-Orange were notably more variable than the others (*ρ*=0.87 and *ρ*=0.80, respectively), and in ways where the two might be mutually compensating for each other’s error when it exists. This may be due to less-than-expected FRET efficiency of the BFP-Orange tandem probe (Fig. S1). All the other probes, however, had quite tight error distributions and fell largely along the x=y line (*ρ*=0.94 to 0.99). These data suggest MuSIC is capable of reasonable quantitative estimation of probe levels from a mixture.

Next, we evaluated agreement between inferred and actual levels by probe and by sample, both quantitatively and with respect to binary classification (Fig. 3F-G). Overall, MuSIC does an excellent job of estimating the presence or absence of probes across samples types, from those containing only two probes, to those containing most or all probes (Fig. 3F). As noted above, the few errors that are noticeable are related to BFP and BFP-Orange (e.g. third 2-way sample from the left), which seem to anti-correlate. One way to evaluate the ability of MuSIC to predict whether a probe is present or absent is by constructing a receiver-operator characteristic (ROC) curve (Fig. 3G). Here, a cutoff for classifying a probe as present or absent is varied, and the performance of classification based on the actual levels is evaluated in terms of true positive and false positive rate. Random classification falls along the x=y dashed line (AUC=0.5). MuSIC has excellent classification performance, identifying nearly all true positives before accumulating false positives (area under the ROC curve = 0.98). Thus, we conclude that MuSIC is capable of both quantitative and binary estimation of at least nine probe levels in a mixture using only ~25% of the available spectrum for excitation. This suggests that future expansion work to the entire spectrum may scale to even greater multiplexing performance. Although we used fluorescent proteins (FPs) here, one can envision mixing and matching both FPs and small molecule fluorophores in a wide potential range of applications, and even bring back in favor fluorophores with complex, multi-modal spectra that may have high information content as a MuSIC probe.

## METHODS

### Computational Methods

#### Availability

All MATLAB code used to perform simulations and analyze data, including the raw data, are included in the supplementary code zip file associated with this manuscript. The scripts “Fig2.m” and “Fig3.m” are the starting points for regenerating the analyses.

#### Data Sources

Data for fluorescent proteins (excitation spectra, emission spectra, brightness) were gathered from individual references (EBFP2^14^, mTagBFP2^15^, mT-Sapphire^16^, mAmetrine^17^, mCerulean3^18^, mTFP1^19^, LSSmOrange^20^, EGFP^21^, TagYFP^22^, mPapaya1^23^, mOrange2^24^, mRuby2^25^, TagRFP-T^24^, mKate2^26^, mCardinal^27^, and iRFP^28^), the Nikon Imaging Center at UCSF (nic.ucsf.edu; fpvis.org), which has subsequently converted to FPbase (fpbase.org), and the Tsien lab website excel file (http://www.tsienlab.ucsd.edu/Documents.htm). All used spectra and other quantitative properties are also contained within the provided MATLAB code.

#### Simulated FRET Efficiency

FRET efficiency *E* is related to the distance between donor and acceptor *r*, and a constant *R_0_* called the Förster radius as follows

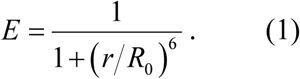

The Förster radius depends on a dipole orientation factor, medium refractive index, donor quantum yield, multiple unit conversion factors and universal constants, and a spectral overlap integral *J*

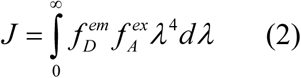

Where *f* is a normalized spectrum (area 1) which depends on wavelength, *D* denotes donor, *A* denotes acceptor, *em* denotes emission, *ex* denotes excitation, and *λ* is wavelength (in microns here for numeric purposes). For the purposes of estimation in the simulation studies, we take all factors except the overlap integral as defined in Eq. 2 as roughly consistent across potential FRET pairs. The overlap integral is calculated using the function `trapz` in MATLAB. We aggregate the constants outside of the overlap integral and estimate its value based on data for mTFP1-mVenus^35^ (73.4, see code). FRET efficiency is capped at 0.35, particularly for the red-shifted proteins which have higher estimated *E* presumably due to the *λ^4^* term in the integral.

#### Simulated Fluorescence Emission Spectra

We calculate the emission intensity spectrum of a probe given a particular excitation wavelength as the sum of multiple emission processes. First, the relative excitation strength of an individual fluorescent protein (FP) is taken from the excitation spectra (with a maximum of 1). Thus, if the excitation wavelength is at the peak of the emission spectra, the excitation strength is 1, and the value along the curve otherwise. The excitation, regardless of the strength, leads to emission distributed across the entire emission spectrum. The overall emission spectra intensity is then multiplied by the relative probe levels (i.e. concentration) and the relative brightness of the FP (comparable across FPs). In the case when a two-FP probe is considered, direct excitation of the donor is calculated as above, but is reduced by the fraction of donor molecules estimated to be undergoing FRET. This fraction is calculated as the estimated FRET efficiency minus the fraction of acceptor that are estimated to be directly excited by the excitation light. We call this the adjusted FRET efficiency. The emission intensity of the acceptor is therefore then composed of two parts scaled by the adjusted FRET efficiency; the first due to FRET from the donor, and the second due to direct acceptor excitation. In the case of a three-FP probe the same logic is extended to the additional interactions to calculate the overall emission spectrum contributions. These calculations are contained within the Supplementary Code in the MATLAB files `CalcIntensity1F.m, CalcIntensity2F.m`, and `CalcIntensity3F.m`. Noise is added to spectra by sampling from a standard normal distribution that is multiplied by 10 (arbitrary units) using the MATLAB function `randn`. This denotes a constant signal to noise of 100 at the maximum probe concentration possible of 1000 (arbitrary units). Thus, this modeled noise has much greater effects on low abundance probes that constitute the majority of the simulated cases.

#### Unmixing

Linear unmixing is performed similarly on simulated and experimental data. First, each emission spectrum is normalized such that the area under the curve is 1 (i.e. divide by sum). For references, the sum used for such normalization is stored for adjustment of probe level estimates later. Then, the MATLAB function `lsqlin` is used to estimate probe levels given a normalized reference matrix and normalized sample vector. Lower bounds of zero and upper bounds of one are used for the levels of each probe, and we constrain the sum of all probe levels to be equal to one, since when the unmixing problem is formulated like this, the probe levels are fractional abundances in the mixture (once corrected for the above noted sums). The function `Unmix.m` in the Supplementary Code contains these calculations. In this case of experimental data, first the averages across triplicates are taken, and then both the reference and mixture data are blank and background (PBS) subtracted prior to analysis.

### Experimental Methods

#### Fluorescent Protein Cloning

Fluorescent protein DNA templates were obtained (LSSmOrange^20^: Addgene #37135; EBFP2^14^: Addgene #14893; ECFP: Gift from Susana Neves; mTFP1 and mVenus, as previously^36^), and sequences amplified by PCR for recombination into DONR221 (Invitrogen) via BP clonase (ThermoFisher Cat: 11789020) following the manufacturer’s protocol. For single probes, a stop codon was included. For dual probes, the first fluorescent protein was amplified without a stop codon, followed at the 3’ end with AgeI-linker-HindIII (linker: GCC GGA GGT GGG GGC CTA GGA). The second fluorescent protein was then added via restriction digest and ligation cloning using the AgeI and HindIII sites. This resulted in an amino acid sequence of TGAGGGGLG between FPs in a tandem probe. Constructs were then recombined using LR Clonase (ThermoFisher Scientific, Cat: 11791100) into pQLinkHD^37^ for protein expression with an N-terminal His-tag (Addgene, Cat: 13668). Constructs were verified by Sanger sequencing.

#### Protein Expression

*E. coli* (BL-21 strain) containing the plasmid-of-interest were taken from a glycerol stock using a sterile pipette tip and added to 5 mL LB (Alfa Aesar, Cat: H26760-36) containing ampicillin (100 ug/mL, Sigma, Cat: A9518-25G) in a 14 mL culture tube (Falcon, Cat: 352063). The cultures were incubated overnight (not more than 16 hours) in an orbital shaker at 37°C and 180 RPM. The following day the culture was added to 100 mL of sterile expression media [1L solution: 20 g bactotryptone (VWR, Cat: 90000-282), 15 g yeast extract (VWR, Cat: 97063-370), 8 g NaCl (Sigma, Cat: S7653-1KG), 4 g NH_2_PO_4_ (Amresco, Cat: 0571-500g), 2 g KH_2_PO_4_ (VWR, Cat: 97062-350) pH to 7.5 with NaOH], 5 mL 40% (40 g / 100 mL) sterile glucose (Alfa Aesar, Cat: A16828) and 100 ug/mL ampicillin in a 1-2L Erlenmeyer flask and incubated in an orbital shaker at 180 RPM and 37°C until OD600 was ~1 (roughly 2-3 hours). Expression was then induced with 1 mM IPTG (VWR, Cat: TCI0328-005G) and incubation continued. For single proteins, cells were cultured overnight at 37°C; for tandem proteins, cells were cultured overnight at 25°C. The next day, cells were pelleted in two 50mL sterile conical tubes (Falcon, Cat: 14-432-22) at 5,000xg for 20 minutes. The pellets were either stored at -20°C until further use or immediately processed for protein purification (see below).

#### Protein Purification

Bacterial cell pellets were resuspended in 15 mL of 1X Wash Buffer [5X Wash Buffer: 250 mM NaH_2_PO_4_ (Amresco, Cat: 0571-500g), 1.5M NaCl (Sigma, Cat: S7653-1KG), 100 mM imidazole (Fisher, Cat: 03196-500), pH adjusted to 8.0 with NaOH]. PMSF (VWR, Cat: 10187-508) was added fresh to each resuspended cell pellet at 1 mM final concentration. The protein solutions were probe sonicated at max power for six cycles of 30 seconds on ice. Subsequently, lysates were spun down at 15,000g for 15 minutes at 4^o^C and the cleared supernatant was collected and kept on ice. A Kimble Flex-Column (VWR, Cat: KT420401-2520) was filled with 25 mL HisPur NiNTA Resin (Thermo Scientific, Cat: 88222), then washed and equilibrated with 15 mL 1X Wash Buffer. The protein supernatant was carefully added to the resin (saving ~500 uL for SDS-PAGE or other analyses later) and allowed to bind to the column for 10-15 minutes. The supernatant was drained from the column and the resin was washed four times with 15 mL 1X Wash Buffer.

After washing, 5 mL of Elution Buffer (50 mM NaH_2_PO_4_ (Amresco, Cat: 0571-500g), 300 mM NaCl (Sigma, Cat: S7653-1KG), 250 mM imidazole (Fisher, Cat: 03196-500), pH 8.0 with NaOH) were added to the column and incubated for 5 minutes. Elution was repeated 10X and each elution was collected. 10 mL PBS (Sigma, Cat: P4417-100TAB) was added to each elution, and then the mixture was added to Amicon Ultra-15 10kDa filter tubes (EMD Millipore, Cat: UFC901024). The tubes were spun at 4000xg for 20 minutes and the flow through was discarded. The concentrate was then washed with 15 mL PBS another 4 times using the same Amicon filter tube. The final concentrate (~200 uL) was kept at 4 °C for further use.

Expression and purification was verified by SDS-PAGE and Coomassie staining of the original lysate supernatant along with all washes and elutions. 50 μL β-mercaptoethanol (Sigma, Cat: M3148-100ML) was added to 950 μL 4X Laemmli buffer (Bio-Rad, Cat: 161-0737) in a microcentrifuge tube. 15 μL of each sample was added to 5 uL of the Laemmli stock in separate 1.5mL microcentrifuge tubes and heated for 5 minutes at 95°C. Mini-Protean TGX Precast Gels (Bio-Rad, Cat: 4569033) with a 15-well comb were used in the BioRad gel electrophoresis cassette (Bio-Rad, Cat: 1658004) with 1L Running Buffer (100 mL 10X Tris/Glycine/SDS Buffer (Bio-Rad, Cat: 1610732) and 900 mL ddH20). Chameleon Precision Plus Protein Dual color standards protein ladder (5 μL; Bio-Rad, Cat:1610374) was used in one lane for each gel, and 15 μL of each sample was added to individual wells. The gels were run at 125V for ~30 minutes until the dye front was close to the bottom. Gels were stained with Coomassie solution [1.2g Coomassie Blue (Thermo Scientific, Cat: 20278) added to 300 mL Methanol (VWR, Cat: BDH1135-4LP) and 60 mL Acetic Acid (Fisher, Cat: A38SI-212)] by microwaving for 30 seconds and then letting the gel stand at room temperature for 10 minutes, and repeating this once. Gels were destained by covering the gel with destaining solution (400 mL methanol (VWR, Cat: BDH1135-4LP) added to 100 ml acetic acid (Fisher, Cat: A38SI-212) and 500 mL of ddH20), microwaving for 45 seconds while covered with saran wrap, adding a few crumbled KimWipes (Fisher, Cat: 06-666A) on top of the gel and allowing the mixture to sit for 10 minutes, discarding the solution, and repeating until the gel was no longer blue. The gel was left in ddH20 overnight and scanned on a LI-COR Odyssey Infrared Scanner at 169 μm resolution and 0.5 mm focus offset for 700-channel fluorescence for visualization. We verified that expressed proteins were the expected molecular weight and the predominant band after purification.

#### Spectral Scanning Measurements

Fluorescence spectra were measured on a Shimadzu RF-5301PC spectrofluorometer. Fluorescent protein probes were diluted with ice-cold PBS (Sigma, Cat: P4417-100TAB) such that peak fluorescence emission intensity was ~1000 AU. Each sample was excited at 10 nm increments from 380 nm to 480 nm. A 2D emission spectral scan was taken of each sample with an emission start 20 nm after the excitation channel center and an emission end at 700 nm. The instrument parameters were set to high sensitivity with an excitation slit width of 5 nm and an emission slit width of 10 nm. Scanning parameters were set to very fast scanning speed, auto response time and 1 nm emission wavelength intervals. Each sample was made in triplicate and individually measured at each excitation wavelength. Data was exported to Excel for further analysis in MATLAB.

## AUTHOR CONTRIBUTIONS

MRB conceived of and supervised the work. HYH, MRB, ADS, MB, CMA, DP and SC performed the experimental work. MRB performed the simulation work. MRB and HYH analyzed data. MRB wrote the paper.

## ACKNOWLEDGEMENTS

MRB acknowledges funding from Mount Sinai, Clemson University and the NIH Grant R21CA196418. MB and ADS were supported by a NIGMS-funded Integrated Pharmacological Sciences Training Program grant (T32GM062754).

---

**Figure S1.**
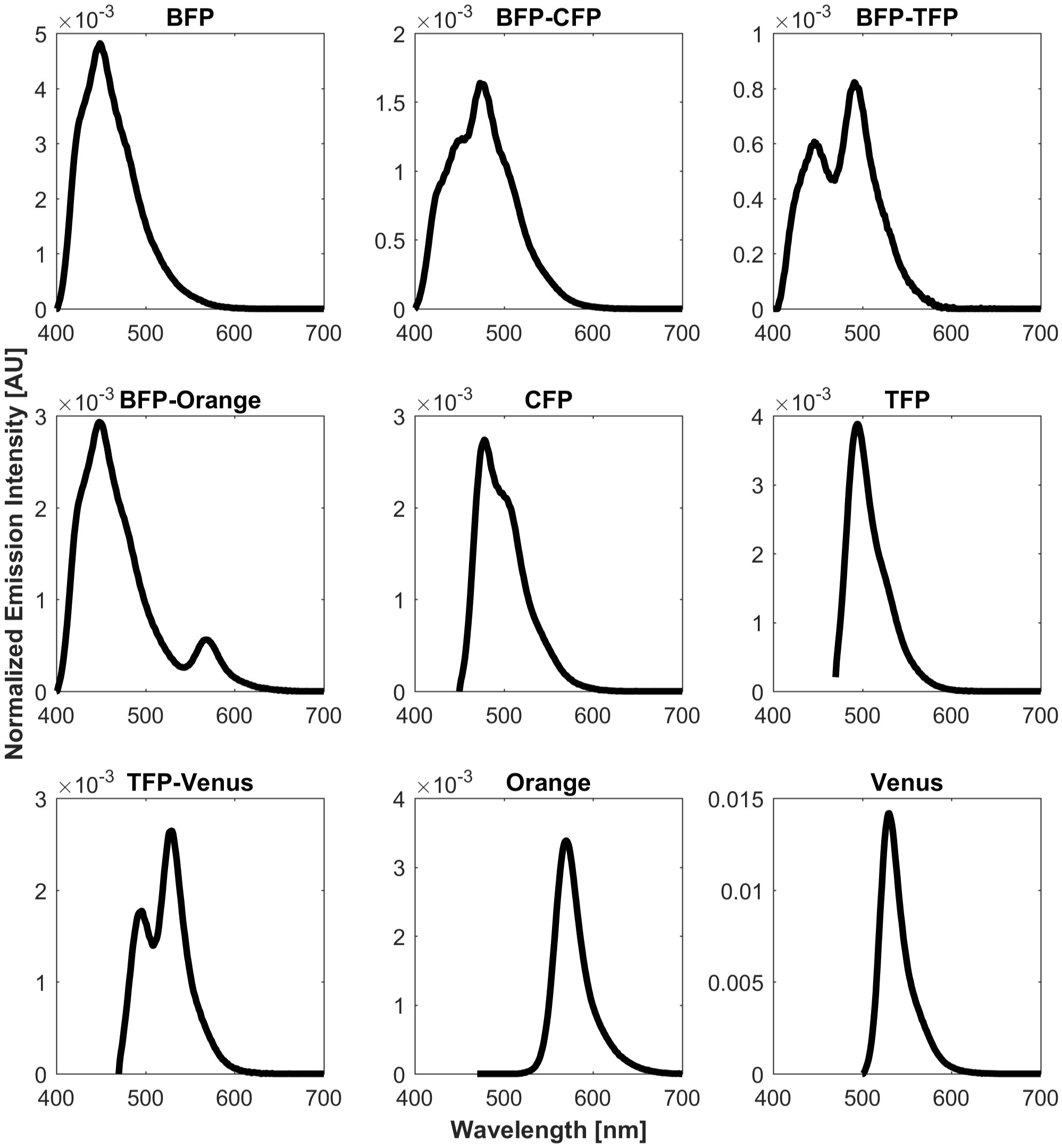
Normalized Emission Spectra of Individual Probes. Emission intensity was background subtracted and then normalized to unit integral. A representative spectra is shown from a suitable excitation channel. BFP, BFP-CFP, BFP-TFP, BFP-Orange: 380nm;CFP: 430 nm; TFP, TFP-Venus, Orange: 450 nm; Venus: 480 nm

